# A multi-omics data simulator for complex disease studies and its application to evaluate multi-omics data analysis methods for disease classification

**DOI:** 10.1101/426510

**Authors:** Ren-Hua Chung, Chen-Yu Kang

## Abstract

An integrative multi-omics analysis approach that combines multiple types of omics data including genomics, epigenomics, transcriptomics, proteomics, metabolomics, and microbiomics, has become increasing popular for understanding the pathophysiology of complex diseases. Although many multi-omics analysis methods have been developed for complex disease studies, there is no simulation tool that simulates multiple types of omics data and models their relationships with disease status. Without such a tool, it is difficult to evaluate the multi-omics analysis methods on the same scale and to estimate the sample size or power when planning a new multi-omics disease study. We developed a multi-omics data simulator OmicsSIMLA, which simulates genomics (i.e., SNPs and copy number variations), epigenomics (i.e., whole-genome bisulphite sequencing), transcriptomics (i.e., RNA-seq), and proteomics (i.e., normalized reverse phase protein array) data at the whole-genome level. Furthermore, the relationships between different types of omics data, such as meQTLs (SNPs influencing methylation), eQTLs (SNPs influencing gene expression), and eQTM (methylation influencing gene expression), were modeled. More importantly, the relationships between these multi-omics data and the disease status were modeled as well. We used OmicsSIMLA to simulate a multi-omics dataset for breast cancer under a hypothetical disease model, and used the data to compare the performance among existing multi-omics analysis methods in terms of disease classification accuracy and run time. Our results demonstrated that complex disease mechanisms can be simulated by OmicsSIMLA, and a random forest-based method showed the highest prediction accuracy when the multi-omics data were properly normalized.

## Introduction

Complex diseases such as hypertension, type 2 diabetes, and autism are caused by multiple genetic and environmental factors (Timpson et al. 2018). Genome-wide association studies have identified many genetic variants (i.e., SNPs) associated with the complex diseases. However, it remains difficult to understand the roles of the associated SNPs in the molecular pathophysiology of the disease and how the SNPs interact with other SNPs in a biological network (Karczewski and Snyder 2018). With the advancement of high-throughput sequencing technology such as next-generation sequencing (NGS) and massive parallel technology such as mass spectrometry, multiple types of omics data (i.e., multi-omics data) including genomics, epigenomics, transcriptomics, proteomics, metabolomics, and microbiomics are rapidly generated (Hasin et al. 2017). As a single type of data generally cannot capture the complexity of molecular events causing the disease, an integrative approach to combining the multi-omics data would be ideal to help elucidate the pathophysiology of the disease (Karczewski and Snyder 2018).

Integrative methods to combine multi-omics data for disease studies have been developed rapidly (Tyekucheva et al. 2011; Jennings et al. 2013; Holzinger et al. 2014; Ruffalo et al. 2015; Yan et al. 2017). They can be generally classified into two categories: multi-staged and meta-dimensional approaches (Ritchie et al. 2015). The multi-staged approach aims to first identify relationships between the multi-omics data, and then test the associations between the multi-omics data and the phenotype. For example, Jennings et al. (Jennings et al. 2013) constructed a Bayesian hierarchical model consisting of two stages. The first stage partitioned gene expression into factors accounted by methylation, copy number variation (CNV), and other unknown causes. These factors were subsequently used as predictors for clinical outcomes in the second stage model. One advantage of this approach is that the causal relationships between multi-omics data can be modeled. In contrast, the meta-dimensional approach combines the multi-omics data simultaneously. Raw or the transformed data from the multi-omics data are combined into a single matrix for the analysis. This approach allows for a more flexible inference of the relationships among the multi-omics data, without the assumptions of the causal relationships between these data.

Although many multi-omics analysis methods for disease studies are available, they were generally evaluated by simulations with data generated specifically to the methods. To compare the performance among these methods, it is necessary to use the same simulated multi-omics dataset with disease status. However, current simulation tools for disease studies mainly focused on simulating a certain type of omics data. For example, more than 25 simulators are available for simulating genetic data with phenotypic trait, according to the Genetic Simulation Resources website (https://popmodels.cancercontrol.cancer.gov/gsr/). Tools such as WGBSSuite (Rackham et al. 2015) and pWGBSSimla (Chung and Kang 2018) can simulate whole-genome bisulphite sequencing (WGBS) data in case-control samples. Moreover, tools such as Polyester (Frazee et al. 2015) and SimSeq (Benidt and Nettleton 2015) simulate RNA-seq data with differential gene expression between two groups of samples. To our knowledge, there is currently no simulation tool that is capable of simulating a variety of omics data types and modeling the complex relationships between the data and the disease. Furthermore, sample size estimation when planning a multi-omics study to ensure sufficient power also becomes important (Hasin et al. 2017). This also requires a simulation tool that simulates realistic multi-omics data structures and models the architecture of the complex disease.

Here, we developed the multi-omics data simulator OmicsSIMLA, which simulates genomics data including SNPs and CNVs, epigenomics data such as the WGBS data, transcriptomics data (i.e., RNA-seq), and proteomics data such as the normalized reverse phase protein array (RPPA) data at a whole-genome level. Furthermore, the relationships between different types of omics data, such as meQTLs (SNPs influencing methylation), eQTLs (SNPs influencing gene expression), and eQTM (methylation influencing gene expression), were modeled. More importantly, the relationships between these multi-omics data and disease status were modeled as well. The disease models in OmicsSIMLA are flexible so that the main effects and/or interaction effects (either risk or protective) of SNPs and CNVs on the disease can be specified. Differential methylation and differential gene and protein expression between cases and controls can also be simulated. We demonstrated the usefulness of OmicsSIMLA by simulating a multi-omics dataset for breast cancer under a hypothetical disease model, and compared the performance among existing multi-omics analysis tools based on the data.

## Results

Figure 1 shows the framework of OmicsSIMLA. The genomics data that can be simulated include SNPs and CNVs. Genotypes at SNPs in unrelated and/or family samples are simulated based on the SeqSIMLA2 algorithm (Chung et al. 2015). CNV status (i.e., a deletion, normal, one duplication or two duplications) on a chromosome is simulated based on the user-specified chromosomal regions and CNV frequencies. Affection status of each sample is determined by a logistic penetrance function conditional on the causal SNPs and CNVs, and/or the interactions among the causal SNPs. The epigenomics data are the methylated and total read counts at CpGs based on bisulphite sequencing, simulated using the pWGBSSimla algorithm incorporating methylation profiles for 29 human cell and tissue types (Chung and Kang 2018). Allele-specific methylation (ASM), in which paternal and maternal alleles have different methylation rates, and differentially methylated region (DMR), where the same CpGs in the region have different methylation rates among different cell types, can also be simulated. Furthermore, the transcriptomics data (i.e., RNA-seq read counts) are simulated with a parametric model assuming a negative-binomial distribution. Finally, the mass-action kinetic action model (Teo et al. 2015) is used to simulate proteomics data at a certain time point incorporating the gene expression data. Some SNPs can be specified as meQTLs and eQTLs, and some CpGs can be specified as eQTM. Allele-specific expression (ASE), which alleles in a gene have different expression levels, caused by cis-eQTL can also be simulated. The differential methylation, gene expression, and protein expression levels between cases and controls are simulated conditional on the affection status.

**Figure 1.**
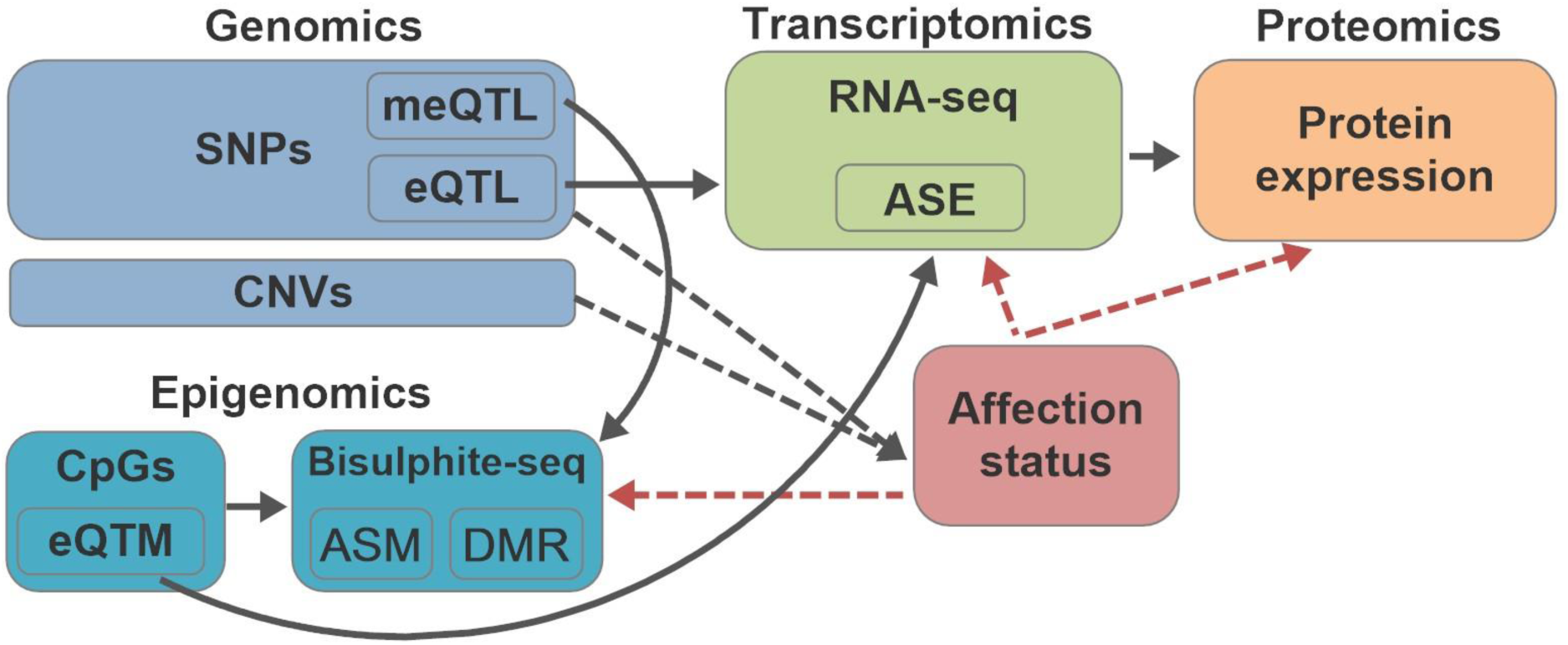
Simulation framework of OmicsSIMLA. The black solid lines represent the relationships among different types of omics data. The black dotted lines represent the causal effects of genomics data to the disease. The red dotted lines represent the retrospective simulations of the methylation, gene expression and protein expression levels conditional on the disease status.

Using OmicsSIMLA, we simulated a multi-omics dataset based on hypothetical pathways for breast cancer as described in Ritchie et al. (Ritchie et al. 2015) and illustrated in Figure 2. The data included a deletion with a protective effect in the CYP1A1 gene, 3 common SNPs with risk effects in the CYP1B1 gene, 5 rare SNPs in the COMT gene, which had interaction effects with a meQTL for the XRCC1 gene, and 5 rare SNPs each in the GSTM1 and GSTT1 genes, which also had interaction effects with eQTLs affecting the gene and protein expression of the XRCC3 gene. A total of 2,022 SNPs in the four genes (i.e., CYP1B1, COMT, GSTM1, and GSTT1) and a regulatory region consisting of the eQTLs and meQTL, 1 CNV in CYP1A1, 688 CpGs in XRCC1, and gene and protein expression levels for 100 genes (including the expression for XRCC3 and 99 other hypothetical genes in the pathways) were simulated. More details about the simulations can be found in the Methods section.

**Figure 2.**
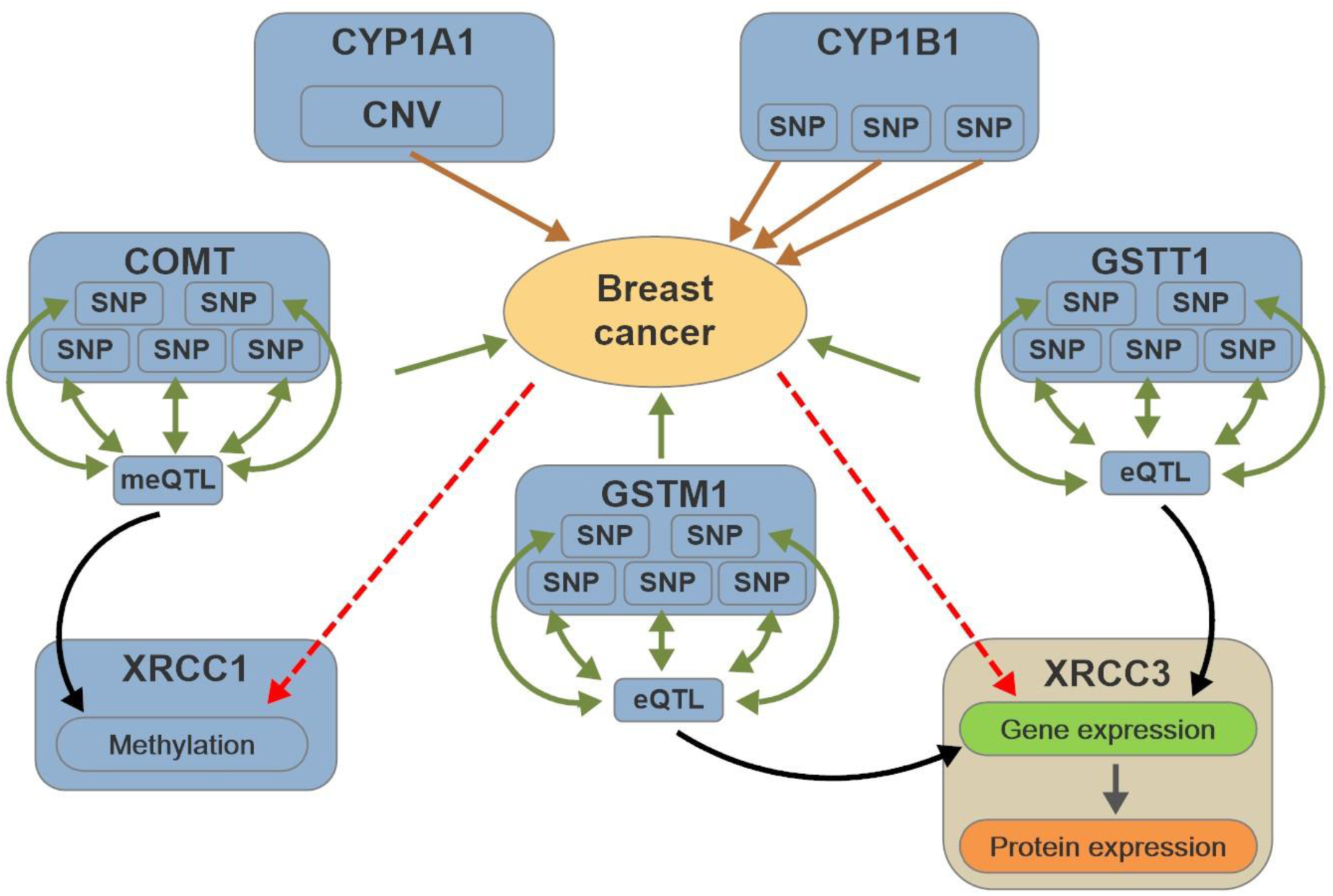
Hypothetical pathways involved in breast cancer. The brown solid lines represent the main effects of SNPs and CNVs on the disease, while the green solid lines represent the interaction effects of SNPs on the disease. The black sold lines represent the regulatory effects of the meQTL and eQTLs on methylation and gene expression, respectively. The red dotted lines represent the retrospective simulations of the methylation, gene expression and protein expression levels conditional on the disease status.

We compared the performance of three multi-omics data analysis methods for disease prediction using the area under the curve (AUC) measures. The three methods included the random forest-based method (RFomics), a graph-based integration method (CANetwork) (Yan et al. 2017), and a model-based integration method (ATHENA) (Holzinger et al. 2014). The RFomics combines the preprocessed multi-omics data in a single matrix for constructing the prediction model. As described in the Methods section, a gene-based risk score is calculated based on SNPs for each gene. Then the risk scores and other multi-omics data are normalized so that they can be evaluated on the same scale by the RF algorithm. In contrast, CANetwork calculates a graph matrix to measure the distance between samples using the composite association network algorithm (Mostafavi et al. 2008), and the prediction model is created based on the distance matrix using the graph-based semi-supervised learning algorithm (Tsuda et al. 2005). Finally, ATHENA creates a neural network model for each type of omics data and a final integrative model is generated based on these models.

Table 1 shows the area under the curve (AUC) for the three methods under three scenarios. Scenario 1 had 500 cases and 500 controls in the training set, and 100 cases and 100 controls in the validation set. Scenario 2 had the same sample sizes as those in Scenario 1, but the multi-omics data had less strong effects on the disease compared to Scenario 1. The effects of the multi-omics data were the same in Scenarios 3 as those in Scenario 1, but Scenarios 3 had larger sample size (i.e., 1,500 cases and 1,500 controls in the training data and 500 cases and 500 controls in the validation data). More details of the three scenarios are provided in the Methods section. Prediction models for the three methods were created based on the training dataset, and their prediction accuracies were evaluated by the validation dataset. As seen in Table 1, RFomics has the highest AUC in all 3 scenarios followed by ATHENA and CANetwork. Table 2 shows the run time for the three methods. In Scenario 1, RFomics and CANetwork had similar performance, while ATHENA required more than 20-times the runtime of RFomics and CANetwork. In Scenario 3, CANetwork was the most efficient method followed by RFomics, and ATHENA also required significantly more time than the other two methods.

**Table 1.** Area under the curve (AUC) for RFomics, CANetwork, and ATHENA under different scenarios

|  | RFomics | CANetwork | ATHENA |
| --- | --- | --- | --- |
| Scenario 1 | 0.861 (0.026)<sup>1</sup> | 0.596 (0.042) | 0.831 (0.042) |
| Scenario 2 | 0.566 (0.041) | 0.529 (0.029) | 0.559 (0.068) |
| Scenario 3 | 0.876 (0.012) | 0.649 (0.019) | 0.835 (0.031) |

<sup>1</sup>The mean AUC and its standard error estimated based on 1,000 batches

**Table 2.** Run time (in seconds) for RFomics, CANetwork, and ATHENA under Scenarios

|  | RFomics | CANetwork | ATHENA |
| --- | --- | --- | --- |
| Scenario 1 | 37.78 | 40.54 | 823.14 |
| Scenario 3 | 143.91 | 94.16 | 2195.95 |

<sup>1</sup>The mean time (in seconds) was estimated based on 100 batches

## Discussion

We have developed OmicsSIMLA, which simulates multi-omics data (i.e., genomics, epigenomics, transcriptomics, and proteomics data) with disease status. In contrast to the current omics data simulators that mainly focused on simulating one type of omics data, OmicsSIMLA simulates multiple types of omics data while the relationships between different types of omics data and the relationships between the omics data and the disease are modeled. As the development of integrative methods for analyzing multi-omics data has attracted substantial interest from researchers, OmicsSIMLA will be very useful to simulate benchmark datasets for comparisons of these methods. Furthermore, as more and more disease studies take advantages of multi-omics data, OmicsSIMLA will also be very useful for power calculations and sample size estimations when planning a new study.

We used OmicsSIMLA to simulate a multi-omics dataset for breast cancer based on hypothetical pathways. Three analysis tools were compared using the dataset. The results suggest that when the different types of data were properly normalized on the same scale, the RF-based method (i.e., RFomics) achieved the highest AUC. Furthermore, RFomics had comparable runtime efficiency as that of CANetwork, while ATHENA was computationally expensive. Therefore, RFomics can potentially be a useful analysis tool for disease prediction using multi-omics data.

Currently, OmicsSIMLA focuses on simulating the dichotomous trait (i.e., affection status). As studies for quantitative traits are also important, it is our future work to extend OmicsSIMLA to simulate quantitative traits based on the classic quantitative genetics model (Falconer and Mackay 1996). Furthermore, environmental factors and the interactions between genes and environments can also play important roles in complex disease etiology. Therefore, simulating exposome data such as the climate and air quality data and modeling their interactions with genes are also important in the future extensions of OmicsSIMLA.

In conclusion, we developed a useful multi-omics data simulator for complex disease studies. As many parameters can be adjusted in OmicsSIMLA, a user-friendly web interface is provided at https://omicssimla.sourceforge.io/generateCommand.html to conveniently specify these parameters.

## Methods

### Simulation of DNA sequences

The SeqSIMLA2 package (Chung et al. 2015) is integrated in OmicsSIMLA to generate DNA sequences in unrelated/related individuals. Similar to SeqSIMLA2, OmicsSIMLA expects a set of external reference sequences (i.e., haplotypes) generated by an external sequence generator, such as COSI (Schaffner et al. 2005) or HAPGEN2 (Su et al. 2011) that has been widely adopted in genetics studies. Generally, a set of 10,000 or more reference sequences are expected. Optional files consisting of recombination rate information and pedigree structures are also accepted. A gene dropping algorithm assuming random mating with crossovers is performed based on the reference sequences, recombination rates, and pedigree structures to generate haplotypes in each individual.

### Simulation of CNVs

For the simulation of CNVs, we considered four CNV states including deletion (D), normal (N), one duplication (U), and two duplications (UU) on a chromosome. Therefore, there are 10 types of CNV states on the two chromosomes in an individual, as shown in Supplemental Table S1, and the total copy numbers on the two chromosomes range from 0 to 6. The user will provide frequencies and ranges of the four CNV states. During meiosis, we use the single-copy crossover model, assuming all crossovers occurred between CNVs (Hartasanchez et al. 2014).

### Simulation of affection status

Genetic variants, including SNPs and CNVs, are used to determine the affection status of an
individual based on a logistic penetrance function as follows:

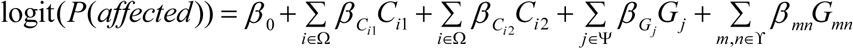

where *P*(*affected*) is the probability of being affected, *β*_0_ determines the baseline prevalence, Ω, Ψ, and ϒ are sets of causal CNVs, SNPs with main effects, and SNPs with interaction effects, respectively, specified by the user, *C_i1_* and *C_i2_* are the CNV states for the first and second haplotypes at CNV *i*, respectively, *G_j_* is the genotype coding at SNP *j*, and *G_mn_* is the genotype coding at SNPs *m* and *n*. *C_i1_ and C_i2_* have values of −1, 0, 1, and 2 for CNV states *D, N, U*, and *UU*, respectively, where *N* is the baseline state. The coding of *G_j_* is based on a dominant, additive or recessive model, and the coding of *G_mn_* is based on several interaction models. If SNP *j* is in a CNV region, allelic CNV (Usher and McCarroll 2015) is considered in the coding of *G_j_*. More details of the coding of *G_j_* and *G_mn_* are provided in the Supplemental Methods. The parameters *β_C_* and *β_G_* are the effect sizes of the main effects for CNVs and SNPs, respectively, and *β_mn_* determines the effect size of the interaction effect between SNPs *m* and *n*. These parameters are specified by the user.

### Simulation of DNA methylation data

The pWGBSSimla package (Chung and Kang 2018) is integrated into OmicsSIMLA to generate the WGBS data. The pWGBSSimla algorithm simulates data using methylation profiles generated based on 41 WGBS datasets for 29 human cell and tissue types. The profiles contain the information for each CpG, such as its distance to the next site, methylation rate, methylation status (i.e., methylated, unmethylated, and fuzzily methylated), and read counts for each type of methylation status. CpGs and the distances between the CpGs are first determined based on the profiles, and then the total read count and methylated read count are simulated for each CpG. Methylation level at a CpG influenced by a meQTL is simulated based on a genotype-specific methylation probability, which is the methylation rate of the CpG in the profiles multiplied by a ratio following an exponential distribution. Furthermore, ASMs are simulated based on father- and mother-specific methylation rates for paternal and maternal alleles, respectively. Finally, a DMR is generated by simulating the same genomic region using profiles for different cell or tissue types. More details of the pWGBSSimla algorithm can be found in Chung and Kang (Chung and Kang 2018).

### Simulation of RNA-seq data

We implemented a parametric simulation procedure for simulating the RNA-seq data similar to that described in Benidt and Nettleton (Benidt and Nettleton 2015). A negative binomial (NB) distribution with mean *μ_ij_* and dispersion parameter *ω_i_* is used to simulate the read count for gene *i* in individual *j*. The mean is calculated as *μ_ij_* = *λ_i_c_j_*, where *λ_i_* is the common mean for gene *i* and *c_j_*. is the individual-specific normalization factor for individual *j*. The parameters ***λ***, ***c***, and *ω* for all genes were estimated using the R package edgeR (Robinson et al. 2010) based on a whole genome RNA-seq dataset consisting of 103 normal tissues in patients with breast cancer from The Cancer Genome Atlas (TCGA) project (Cancer Genome Atlas Research 2008). The parameters *λ_i_* and *ω_i_* are randomly sampled with replacement from ***λ*** and ***ω***. If more than 103 samples are simulated, we use the smoothed bootstrap procedure (Efron and Tibshirani 1993) to calculate 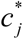 for individual *j*, and *μ_ij_* is calculated as *λ_i_*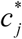. More details of the calculation of 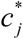 are provided in the Supplemental Methods. The user can specify *n* differentially expressed (DE) genes between cases and controls and their fold changes, and the read count for DE gene *i* in individual *j* is simulated based on a NB distribution with mean *f_i_μ_ij_* and dispersion parameter *λ_i_*, where *f_i_* is the fold change for gene *i*.

### Simulation of eQTL and allele-specific reads

We followed the procedure in the simulation study in Sun (Sun 2012) to simulate eQTL and read counts for ASE. For eQTL *l* with a user-specified fold change *hl*, the means for the three genotypes *AA*, *Aa*, and *aa* at the eQTL are *μ_ij_*, *h_l_μ_ij_*, and (2*h_l_* – 1)*μ_ij_*, respectively, and the dispersion parameter is *ω_i_* in the NB distribution for gene *i* influenced by the eQTL. ASE for a gene caused by a cis-eQTL is simulated by assuming reads were mapped to heterozygous SNPs (i.e., allele-specific reads) in the gene. A cis-eQTL refers to the eQTL being located in the cis-regulatory elements of the gene. Because the alleles at the cis-eQTL can be in the same haplotype as the alleles of the gene, ASE can be observed using the allele-specific reads of the gene. Furthermore, only heterozygous SNPs can be tested for cis-eQTL with the allele-specific reads. Therefore, we simulate allele-specific reads for heterozygous eQTLs. Assuming *t_ij_* is the total read count for gene *i* in individual *j*, the total number of allele-specific reads is calculated as 0.005*t_ij_* where 0.005 was estimated from real data by Sun (Sun 2012). Furthermore, also suggested by Sun (Sun 2012), the number of allele-specific reads for a haplotype is simulated using a beta-binomial distribution with a mean determined by the effect size of the cis-eQTL and an overdispersion parameter of 0.1. The effect size is defined as log_2_(expression of the alternative allele at the eQTL/expression of the reference allele at the eQTL) (Mohammadi et al. 2017) for a heterozygous cis-eQTL and is set to 0 for a homozygous cis-eQTL.

### Simulation of eQTM

We used linear regression to model the relationship between gene expression and methylation: 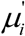 = *E(y_ij_)* = *α_i_* + *β_ij_*, where *y_ij_* and *x_ij_* are the RNA-seq read count and the proportion of methylated reads, respectively, for gene *i* influenced by methylation in individual *j*. Assuming that the NB parameters for gene *i* are *μ_i_* and *ϕ_i_*, the parameter *α_i_* is specified as *μ_i_*, and *β_i_* is assumed to follow a normal distribution with a mean and a standard deviation specified by the user. Then the gene expression of gene *i* is simulated by an NB distribution with parameters of 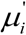 and *ϕ_i_*.

### Protein expression simulation

We assumed that the protein expression level for protein *k* at a time point *t* in sample *j* follows a normal distribution with a mean *η_kjt_* and a standard deviation *τ_k_* after normalization. We used the mass-action kinetic action model (Teo et al. 2015) to simulate protein expression at a certain time point. The mean *η*_*kj*,*t*+1_ for the protein expression at time *t*+1 was determined as follows:

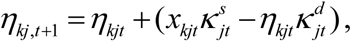

where *x_kjt_* is the normalized gene expression for the gene encoding protein *k*, and 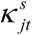 and 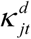 are the protein synthesis and degradation rates, respectively, in individual *j* at time *t*. The normalized gene expression *x_kjt_* is calculated using the median absolute deviation (MAD) scale normalization (Fundel et al. 2008) based on the RNA-seq data simulated from the previous section. Similar to the simulation study in Teo et al. (Teo et al. 2015), 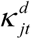 is fixed to be 1, and 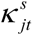 with a default value of 1 can be changed by the user. A vector of standard deviations ***τ*** were estimated from the level 4 protein expression data of primary tumor tissue in 874 breast cancer patients from the TCGA project downloaded from the cancer proteome atlas (TCPA) (Li et al. 2013) website. The level 4 data consist of protein expression data for 224 proteins that have been normalized across the samples as well as across the proteins, and a replication-based method was used to account for differences in protein expression among different batches. The parameter *τ_j_* is then randomly sampled with replacement from ***τ***.

### A random-forest based method for integrating multi-omics data for disease studies

Multi-omics data can have different data types (e.g., discrete data for SNP genotypes, categorical data for CNV statuses, and continuous data for proportions of methylated reads, RNA-seq read counts, and normalized protein expression) and different variations (e.g., three possible values of 0, 1, and 2 for minor allele counts at SNPs, and real numbers ranging between 0 and 1 for the proportions of methylated reads). When developing a method for integrating these data, it is important to account for the properties of different data types so that the analysis results would not be biased toward certain variables (Ritchie et al. 2015). We developed a preprocessing algorithm for the multi-omics data. A gene-based risk score, which is a weighted sum of the numbers of risk alleles at SNPs in the gene, for each individual is constructed. The weights are the effect sizes of the risk alleles at the SNPs. More details for calculating the risk score are provided in the Supplemental Methods. Then each variable from different omics data, including the gene-based risk scores, CNV statuses of genes, methylation proportions at CpGs, gene and protein expression levels, is normalized so that it has a mean 0 and a standard deviation of 1. The normalized variables are then used in RF for classification.

### Simulation studies

We used OmicsSIMLA to evaluate the performance of the proposed RF-based method, compared with CANetwork and ATHENA. A hypothetical disease model for breast cancer involving multi-omics data (Ritchie et al. 2015) was simulated, as shown in Figure 2. To be more specific, a deletion with a frequency of 20%, which had a protective effect with an odds ratio (OR) of 0.67, in the CYP1A1 gene and 3 common variants, which had main effects (ORs = 1.5) with minor allele frequencies (MAFs) > 10%, in the CYP1B1 gene were simulated. We also simulated 5 rare variants with MAFs < 3% in the COMT gene, which had interaction effects (ORs = 5) with a meQTL for the XRCC1 gene. The CpG in XRCC1 influenced by the meQTL caused a difference in methylation rates of 10% between cases and controls. Furthermore, we simulated 5 rare variants in the GSTM1 gene, which had interaction effects (ORs = 5) with a cis-eQTL for the XRCC3 gene, and 5 rare variants in the
GSTT1 gene, which had interaction effects (ORs = 5) with a trans-eQTL for XRCC3. The eQTL caused a fold change of 1.5 in the XRCC3 gene expression compared to the reference genotype, and a fold change of 1.5 was simulated for the differential gene expression of XRCC3 between cases and controls. In summary, the total variables consisted of 200, 687, 264, and 176 SNPs in the CYP1B1, COMT, GSTM1, and GSTT1 genes, respectively, and 695 SNPs harboring the meQTL and two eQTLs in the regulatory region, a variable for CNV status in CYP1A1, methylation levels at 688 CpGs in XRCC1, and gene and protein expression levels for 100 genes and their encoded proteins. More details for generating the reference sequences in the genes and the simulations for each omics data type are provided in the Supplemental Methods.

We simulated a training dataset consisting of 500 cases and 500 controls as well as a validation dataset consisting of 100 cases and 100 controls. The training dataset was used by RFomics, CANetwork, or ATHENA to construct a prediction model. The validation dataset was then used to calculate the AUC based on the prediction model. Note that a 5-fold cross-validation was performed in ATHENA, and a best model based on the testing dataset (i.e., one of the five random 20% of the training dataset) was created for each cross-validation. The model with the highest AUC based on the testing dataset was selected and applied to the validation dataset. This simulation scenario was referred to as Scenario 1. We also simulated a scenario with less strong genetic effects (Scenario 2) and a scenario with larger sample size (Scenario 3). More details about Scenarios 2 and 3 are provided in the Supplemental Methods. For each scenario, 1,000 batches of training and validation datasets were simulated, and the AUC for each algorithm was averaged over the 1,000 batches.

## Data access

The simulated data generated in the simulation studies can be downloaded from the OmicsSIMLA website (https://omicssimla.sourceforge.io/download.html).

## Acknowledgements

This work has been supported by a grant from the Ministry of Science and Technology (MOST 106-2221-E-400-005-MY3) in Taiwan.

## References

Benidt S, Nettleton D. 2015. SimSeq: a nonparametric approach to simulation of RNA-sequence datasets. Bioinformatics 31: 2131–2140.

Cancer Genome Atlas Research N. 2008. Comprehensive genomic characterization defines human glioblastoma genes and core pathways. Nature 455: 1061–1068.

Chung R-H, Kang C-Y. 2018. pWGBSSimla: a profile-based whole-genome bisulphite sequencing data simulator incorporating methylation QTLs, allele-specific methylations and differentially methylated regions. bioRxiv doi:10.1101/390633.

Chung RH, Tsai WY, Hsieh CH, Hung KY, Hsiung CA, Hauser ER. 2015. SeqSIMLA2: simulating correlated quantitative traits accounting for shared environmental effects in user-specified pedigree structure. Genetic epidemiology 39: 20–24.

Efron B, Tibshirani RJ. 1993. An Introduction to the Bootstrap. Chapman and Hall/CRC.

Falconer DS, Mackay TF. 1996. Quantitative genetics. Benjamin Cummings, San Francisco.

Frazee AC, Jaffe AE, Langmead B, Leek JT. 2015. Polyester: simulating RNA-seq datasets with differential transcript expression. Bioinformatics 31: 2778–2784.

Fundel K, Kuffner R, Aigner T, Zimmer R. 2008. Normalization and gene p-value estimation: issues in microarray data processing. Bioinform Biol Insights 2: 291–305.

Hartasanchez DA, Valles-Codina O, Braso-Vives M, Navarro A. 2014. Interplay of interlocus gene conversion and crossover in segmental duplications under a neutral scenario. G3 4: 1479–1489.

Hasin Y, Seldin M, Lusis A. 2017. Multi-omics approaches to disease. Genome biology 18: 83.

Holzinger ER, Dudek SM, Frase AT, Pendergrass SA, Ritchie MD. 2014. ATHENA: the analysis tool for heritable and environmental network associations. Bioinformatics 30: 698–705.

Jennings EM, Morris JS, Carroll RJ, Manyam GC, Baladandayuthapani V. 2013. Bayesian methods for expression-based integration of various types of genomics data. EURASIP J Bioinform Syst Biol 2013: 13.

Karczewski KJ, Snyder MP. 2018. Integrative omics for health and disease. Nature reviews Genetics 19: 299–310.

Li J, Lu Y, Akbani R, Ju Z, Roebuck PL, Liu W, Yang JY, Broom BM, Verhaak RG, Kane DW et al. 2013. TCPA: a resource for cancer functional proteomics data. Nature methods 10: 1046–1047.

Mohammadi P, Castel SE, Brown AA, Lappalainen T. 2017. Quantifying the regulatory effect size of cis-acting genetic variation using allelic fold change. Genome research 27: 1872–1884.

Mostafavi S, Ray D, Warde-Farley D, Grouios C, Morris Q. 2008. GeneMANIA: a real-time multiple association network integration algorithm for predicting gene function. Genome biology 9 Suppl 1: S4.

Rackham OJ, Dellaportas P, Petretto E, Bottolo L. 2015. WGBSSuite: simulating whole-genome bisulphite sequencing data and benchmarking differential DNA methylation analysis tools. Bioinformatics 31: 2371–2373.

Ritchie MD, Holzinger ER, Li R, Pendergrass SA, Kim D. 2015. Methods of integrating data to uncover genotype-phenotype interactions. Nature reviews Genetics 16: 85–97.

Robinson MD, McCarthy DJ, Smyth GK. 2010. edgeR: a Bioconductor package for differential expression analysis of digital gene expression data. Bioinformatics 26: 139–140.

Ruffalo M, Koyuturk M, Sharan R. 2015. Network-Based Integration of Disparate Omic Data To Identify “Silent Players” in Cancer. PLoS computational biology 11: e1004595.

Schaffner SF, Foo C, Gabriel S, Reich D, Daly MJ, Altshuler D. 2005. Calibrating a coalescent simulation of human genome sequence variation. Genome research 15: 1576–1583.

Su Z, Marchini J, Donnelly P. 2011. HAPGEN2: simulation of multiple disease SNPs. Bioinformatics 27: 2304–2305.

Sun W. 2012. A statistical framework for eQTL mapping using RNA-seq data. Biometrics 68: 1–11.

Teo G, Vogel C, Ghosh D, Kim S, Choi H. 2015. A Mass-Action-Based Model for Gene Expression Regulation in Dynamic Systems. Cambridge University Press.

Timpson NJ, Greenwood CMT, Soranzo N, Lawson DJ, Richards JB. 2018. Genetic architecture: the shape of the genetic contribution to human traits and disease. Nature reviews Genetics 19: 110–124.

Tsuda K, Shin H, Scholkopf B. 2005. Fast protein classification with multiple networks. Bioinformatics 21 Suppl 2: ii59–65.

Tyekucheva S, Marchionni L, Karchin R, Parmigiani G. 2011. Integrating diverse genomic data using gene sets. Genome biology 12: R105.

Usher CL, McCarroll SA. 2015. Complex and multi-allelic copy number variation in human disease. Briefings in functional genomics 14: 329–338.

Yan KK, Zhao H, Pang H. 2017. A comparison of graph- and kernel-based-omics data integration algorithms for classifying complex traits. BMC bioinformatics 18: 539.

